# Genomic 8-oxoguanine modulates gene transcription independent of its repair by DNA glycosylases OGG1 and MUTYH

**DOI:** 10.1101/2024.02.20.581185

**Authors:** Tobias Obermann, Teri Sakshaug, Vishnu Vignesh Kanagaraj, Andreas Abentung, Antonio Sarno, Magnar Bjørås, Katja Scheffler

## Abstract

8-oxo-7,8-dihydroguanine (OG) is one of the most abundant oxidative lesions in the genome and associated with genome instability. Its mutagenic potential is counteracted by a concerted action of 8-oxoguanine DNA glycosylase (OGG1) and mutY homolog DNA glycosylase (MUTYH). It has been suggested that OG and its repair has epigenetic-like properties and mediates transcription, but genome-wide evidence of this interdependence is lacking. Here, we applied an improved OG-sequencing approach reducing artificial background oxidation and RNA-sequencing to correlate genome-wide distribution of OG with gene transcription in OGG1 and/or MUTYH-deficient cells. Our data identified moderate enrichment of OG in the genome that is mainly dependent on the genomic context and not affected by DNA glycosylase-initiated repair. Interestingly, no association was found between genomic OG deposition and gene expression changes upon loss of OGG1 and MUTYH. Regardless of DNA glycosylase activity, OG in promoter regions correlated with expression of genes related to metabolic processes and damage response pathways indicating that OG functions as a cellular stress sensor to regulate transcription. Our work provides novel insights into the mechanism underlying transcriptional regulation by OG and DNA glycosylases OGG1 and MUTYH and suggests that oxidative DNA damage accumulation and its repair utilize different pathways.

## INTRODUCTION

Reactive oxygen species (ROS) are one of the major threats for DNA integrity (1). ROS leads to accumulation of oxidative DNA damage which is a potential source of mutagenesis, which may lead to cancer, aging or neurodegenerative diseases (2,3). Due to its low redox potential, the DNA base guanine is most prone to oxidation, resulting in, among others 8-oxo-7,8-dihydroguanine (OG) which is estimated to occur on average about 100 to 500 times per cell each day (4). OG is recognized and excised by 8-oxoguanine DNA glycosylase (OGG1) through a process called the base excision repair (BER) pathway (5). OG is a pre-mutagenic lesion, as during replication adenine can be incorporated opposite of OG, giving rise to a stable and pro-mutagenic OG:A base pairing (6). A second round of replication results in a T:A base pair manifesting the oxidative damage into a point mutation (7). DNA glycosylase mutY homolog (MUTYH) repairs the misincorporated adenine opposite of OG thereby preventing mutations (6). Previously we have shown that global levels of OG are not increased in the brain of mice lacking OGG1 and/or MUTYH but that their hippocampal transcriptome is altered (3). Differentially expressed genes overlapped between the different knockout (KO) mice and were associated with anxiety, learning and memory pathways (3). However, the molecular mechanism behind this remains mainly unknown.

Several recent studies suggested a role for OG as an epigenetic-like mark. Stress-induced accumulation of OG and recognition by OGG1 in promoter regions was shown to regulate gene expression of specific genes (8,9). In addition, apurinic (AP) sites generated by excision of OG activates gene transcription, which has been demonstrated for vascular endothelial growth factor (*VEGF*), endonuclease III-like protein 1 (*NTHL1*) (10) and sirtuin 1 (*SIRT1*) genes (11). Furthermore, OG has been shown to destabilize G-quadruplex (G4) formations in telomeric regions (12). This secondary DNA structure consists largely of Gs and is involved in the regulation of transcription (13,14). Upon excision of OG on the coding strand in potential G4 forming sequences, the resulting abasic site and subsequent AP endonuclease1 (APE1) binding enabled G4 formation and upregulation of gene transcription of the VEGF gene (10,15). Alternatively, within the Kirsten ras (*KRAS*) gene, OG was formed under oxidative stress within the folded G4 causing the recruitment of nuclear factors that unfolded the secondary G4 structure (16). This enabled OGG1 to repair OG and subsequently transcriptional upregulation of *KRAS* was observed.

To better address the role of OG in genome regulation several OG-sequencing (OG-seq) methods have been established differing mainly in the mechanisms of OG recognition (anti-OG antibody (17), selective chemical reaction (18), OG-specific glycosylases (19–21) or chemical labeling and polymerase stalling (22)). Together, these studies demonstrated a non-random genomic distribution of OG with regulatory, CG-rich and G4 regions being mostly affected. However, conclusions are inconsistent and might be driven by artificial introduction of OG during sample preparation or inadequate data analysis approaches (1). So far only one study directly addressed the consequence of deficient repair on genome-wide OG damage distribution and showed increased OG level particularly in promoter, 5‘ untranslated regions (UTRs) and 3‘ UTRs in OGG1-deficient mouse embryonic fibroblasts (18).

In the present study, we investigate the genome-wide distribution of OG in HAP1 cells deficient for OGG1 and/or MUTYH by using an improved OG-seq approach that reduces artificial background oxidation and compared it to gene expression levels using RNA-sequencing. This allows for the first time to address the suggested epigenetic-like properties of OG and its repair in transcriptional regulation on a genome-wide scale *in vivo*.

## MATERIAL AND METHODS

### Cell culture

The near-haploid HAP1 cells are derived from the KBM-7 cell line and were obtained from Horizon Discovery. The cells were cultured in Iscove’s Modified Dulbecco’s Medium with 10% Fetal Bovine Serum and 1% Penicillin/Streptomycin. The insertions/deletions generated by CRISPR-Cas9 technology in the *OGG1* and *MUTYH* genes are provided in Supplementary Table 1. Two clones of each cell line were used for the experiments if not indicated otherwise.

### Proliferation and survival assay

HAP1 cells were seeded in triplicate onto a 96-well plate at a concentration of 5 000 cells/well. For proliferation, cells were grown for 24 h-72 h. For cell survival, cells were treated 24 h after seeding with different concentrations of hydrogen peroxide for 24 h. Viability was measured using PrestoBlue cell viability reagent according to the manufacturers protocol. Fluorescence was measured using FLUOstar Omega Plate reader (BMG Labtech).

### Nucleic acid isolation

Genomic DNA was isolated using the DNeasy Blood & Tissue Kit (Qiagen) and total RNA was isolated using the RNeasy Kit (Qiagen) according to the manufactures protocol. To prevent DNA oxidation 0.1 mM Butylated hydroxytoluene (BHT) and 0.1 mM Deferoxamine (DFO) were added during the isolation procedure. DNA and RNA concentrations were measured using NanoDrop (Thermo Fisher).

### Global OG measurement

Global OG measurements were performed on genomic DNA and fragmented DNA. Prior to hydrolysis, fragmented DNA samples were purified by isopropanol precipitation. 0.3 volumes of 10 M ammonium acetate and 1 volume of isopropanol were added to the DNA samples. The samples were then centrifuged at 16 000 g for at least 30 min. Next, the supernatants were discarded by decanting and samples were washed twice with ice-cold 70% ethanol. Finally, the pellets were dried at room temperature for 5 min.

To hydrolyze DNA, samples were incubated for 1 h at 37 °C in DNA hydrolysis mixture consisting of hydrolysis buffer (10 mM NH_4_Ac, 1 mM MgCl_2_, 0.1 mM ZnCl_2_) with 0.008 U/μl nuclease P1 (NP1), 0.8 U/μl benzonase, and 0.04 U/μl alkaline phosphatase, including the antioxidants BHT and DFO with a final concentration of 0.1 mM each. The reaction was stopped by placing the samples on ice. Next, proteins were precipitated from the solution by adding 150 μl ice-cold acetonitrile followed by 30 min centrifugation at 4 °C 16 000 g. Subsequently, the supernatant was transferred into new tubes and lyophilized at - 80 °C overnight. Finally, the samples were dissolved in water. Single nucleotide 8-oxoguanine (O(dG)) was analyzed by liquid chromatography-mass spectrometry (LCMS) using an Agilent 6495 triple quadrupole LCMS system with an Agilent EclipsePlusC18 RRHD column (2.1 x 150 mm, 1.8 µm particle size) at 25 °C. The mass transition used was 284.1 à 168.0 m/z. The mobile phases were (A) UHPLC-grade water and (B) UHPLC-grade methanol, both containing 0.1% UHPLC-grade formic acid. The HPLC method used a flow rate of 230 ml/min with 5% B to 0.5 min, ramp to 20% B at 2.5 min, ramp to 95% B at 6 min, hold at 95% B from 6 to 7 min, ramp to 5% at 7.2 min, and equilibration with 5% B from 7.2 to 10.5 min. For unmodified dNs, we diluted samples 1:5000 and analyzed them on a API5000 triple quadrupole mass spectrometer (Applied Biosystems) with an Acentis® Express C18 column (0.5 x 150 mm, 2.7 µm particle size) at 40 °C. The HPLC method used a flow rate of 200 µl/min with an isocratic flow of 22% B for 3 min. The mass transitions used were 252.1 à 136, 228.1 à 111.9, 268.1 à 152, and 243.1 à 127 m/z for dA, dC, dG, and dT, respectively.

### RNA-sequencing

Total RNA from two clones of each genotype was sent to BGI Tech Solutions Co., Hong Kong for RNA-sequencing on DNBseq platform. FASTQ files were quality assessed using FastQC and Fastq-Screen (https://www.bioinformatics.babraham.ac.uk/projects/) and aligned to the GRCh38 genome using the splice-aware aligner Hisat2 (v.2.2.1; (23)). All samples had over 97% alignment rate. Quantification of reads over genes was performed by the Bioconductor Rsubread (v2.2.6) package in R (v4.2.2) using annotations from the Ensembl GRCh38.p13 (release 103) reference. Normalization and differential expression analysis was then performed between genotypes using the quasi-likelihood method from the Biocondictor package edgeR (v.3.42.4; (24)). Genes were considered differentially expressed if they had a log2FC beyond ± 1 and a P. value ≤ 0.05. Volcano plots for RNA-seq data were generated using VolcanosR2 (https://huygens.science.uva.nl/VolcaNoseR2/ https://huygens.science.uva.nl/VolcaNoseR2/). Pathway analysis was performed on differentially expressed genes (DEGs) for gene ontology (GO) biological pathways using Pantherdb overrepresentation fisher test and p-values were corrected for false discovery rate. A normalized count matrix was exported from edgeR for further analysis and comparison.

### DNA Fragmentation

Antioxidants BHT and DFO were added with a final concentration of 0.1 mM to the samples during fragmentation to avoid artificial introduction of OG. For the sonication procedures, genomic DNA was prepared in 10 mM Tris-HCl (pH 8.0), 100 μM DFO, and 100 μM BHT. The sonicators were operated according to the manufacturers’ protocols to obtain a target fragment size of 150bp. Specifically, the Diagenode sonicator was operated in 30 cycles, wherein one cycle consists of 30 s on and 30 s off. The Covaris sonicator was operated with 50 W peak incident power, a duty factor of 20%, 200 cycles per burst and with a total treatment time of 350 s. DNA fragmentation by Fragmentase was performed using 3 µg genomic DNA in the supplied 1x Fragmentase buffer (20 mM Tris-HCl, 15 mM MgCl_2_, 50 mM NaCl, 0.1 mg/ml bovine serum albumin, 0.15% Triton X-100, pH 7.5) and additional 0.1 mM of BHT and DFO. After 30 min at 37 °C the reaction was stopped by placing the samples on ice and by adding 5 µl ice-cold EDTA. Fragmented DNA was cleaned up using DNA Clean & Concentrator-5 (Zymo Research). To visualize the fragmentation, DNA was loaded onto 2% agarose gel including 1X GelRed (Biotium).

### OG-Sequencing

Affinity purification of OG containing DNA fragments was performed according to Ding et al. (18) with minor modifications. In brief, fragments were chemically labeled using 20 mM amine-terminated biotin with a polyethylene glycol linker (BTN) and 5 mM K_2_IrBr_6_ in 100 mM NaP_i_ (pH 8.0) for 30 min at 45 °C. Subsequently, a PCR purification kit (Qiagen) was used to remove excess BTN. Labeled fragments were purified using Dynabeads MyOne Streptavidin C1 (Invitrogen) washed and eluted by incubation in 150 mM NaOH for 30 min. Prior to library preparation single stranded (ss) DNA was concentrated using the ssDNA/RNA clean & concentrator kit (ZYMO).

Libraries were constructed using the Accel-NGS® 1S Plus DNA Library Kit (Swift Biosciences™) according to the protocol with an initial input of 500 pg per sample. Electropherograms (High Sensitivity DNA Assay, Bioanalyzer, Agilent Technologies) were performed prior to pooling. Pooled libraries from two clones of each genotype (in total three independent experiments) were sent to BGI Tech Solutions Co., Hong Kong for sequencing on a MGISEQ-2000 platform. The samples were sequenced as paired end reads of 100bps.

Illumina adapters were trimmed from raw files using cutadapt (v.2.8) and quality of the reads was assessed using FastQC and Fastq-Screen. Datasets were mapped from FASTQ files to the human genome (GRCh38) with Bowtie2 (v.2.3.5.1; (25)) using default settings and duplicate reads were marked using samtools (v1.10). Reads were counted over standard chromosomes (i.e. chr 1-22 & X) in non-overlapping 1000bp bins. Incomplete bins (i.e. smaller than 1000bp) or bins overlapping blacklisted regions (26) were removed. Reads were counted using bedtools (multicov; v2.27.1) and counts were normalized using a rank statistics-based signal extraction scaling (SES) method as previously published by Diaz et al. (27). This method works by differentiating between genomic regions with enrichment in a targeted pull-down sample and background regions found in both the pull-down and the input to use only background regions for normalization. Briefly, binned counts are reordered by OG signal intensity and the difference of the cumulative percentage tag by ordered bin was calculated to differentiate signal from noise regions. An inter-library scaling factor was then calculated from only low-damage background regions, which was applied to input counts. Genotypes showed no difference in global accumulation of oxidative damage (Figure 1C), but it was important to consider that differences in library size between genotypes at specific genomic elements could be dependent on genotype. Therefore, inter-genotype library scaling factors used in all regional analyses were calculated from total library size as the ratio of the individual sample to the mean of all samples over all bins.

**Figure 1:**
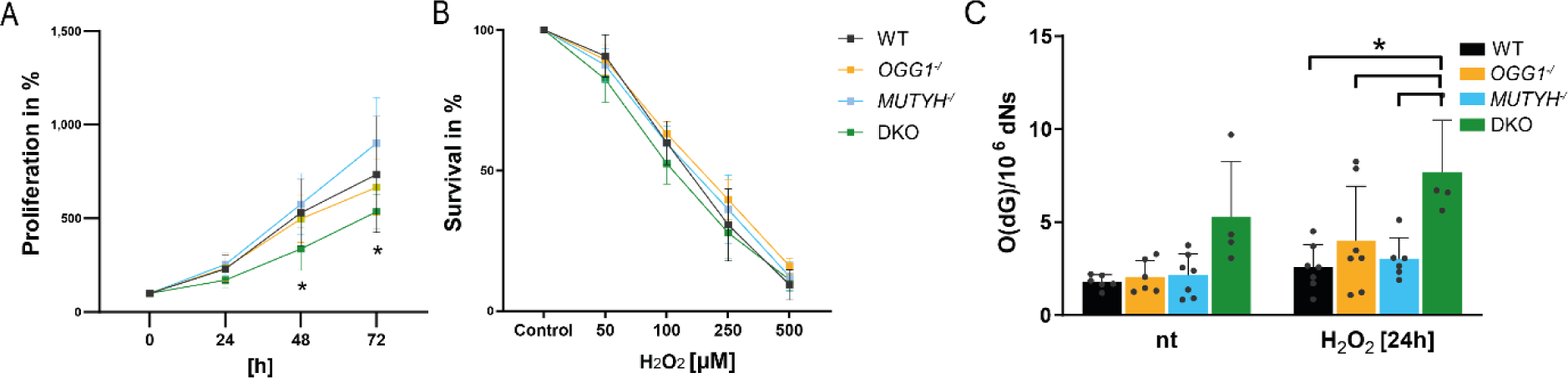
Characterization of WT and DNA glycosylase deficient HAP1 cells. **(A)** Proliferation of WT, *OGG1^-/^*, *MUTYH^-/^* and DKO over the course of 72 h. **(B)** Survival upon treatment with increasing concentrations of H_2_O_2_. **(C)** Global levels of single nucleotide 8-oxoguanine (O(dG)) of untreated (nt) cells and 24 h after H_2_O_2_ treatment. Statistical analyses were done in GraphPad Prism software 10.1 using 2-way ANOVA with Dunnett’s multiple comparisons test. P-values < 0.05 were considered significant. Error bars represent standard deviation from the mean based on three independent experiments and two clones per genotype except for DKO in (C) where only one clone was used.

### Enrichment of OG across the genome

Areas of enrichment were identified by the log2 ratio of OG to Input signals over 1000bp bins. To ensure biological relevance, technical replicates were processed individually and only bins with the largest log2 fold-change were considered (≥ 95^th^ percentile). Extracted bins were compared and retained only if they were present in both technical replicates and at least two of three biological replicates. Bookended bins were combined within genotype and annotated using Homer (v4.11.1) to closest gene and genomic feature. GC content of enriched regions was determined using bedtools.

### Enrichment of OG at genomic features

To assess the difference in OG accumulation across different genomic features, reads were analyzed over the following regions: 1000bp upstream of gene start, CpG islands, exons, introns, 1000bp downstream of gene end and intergenic regions. Regional bed files were downloaded for the hg38 genome from the UCSC table browser (http://genome.ucsc.edu/cgi-bin/hgTables) for CpG regions, exons, introns, upstream, downstream, and whole genes. Intergenic regions were determined by taking the complement of whole genes over the genome using bedtools. Blacklisted regions were subtracted from all regional bed files. Samples were counted over regions individually and imported into R for processing. Input counts were normalized using signal extraction scaling (SES) factors and subtracted from OG reads to a minimum of 0. To find genotype-specific enrichment of oxidative damage, low-read filtering and differential enrichment was performed in edgeR using manually set predetermined library normalization factors. Differentially enriched regions were defined as all regions with a P. value ≤ 0.01 and a log2FC beyond ± 0.5. Results were annotated using Homer and gene lists from differentially enriched CpG islands, exons, introns and 1000bp upstream of gene start were analyzed using Pantherdb for overrepresentation of genes in GO biological processes.

### Enrichment of OG in G4 motifs

G4 regions were defined using DeepG4 (28). DeepG4 uses a deep-learning approach to score DNA regions on their potential to form G4 complexes based on sequence motifs. Here we used DeepG4 to scan the hg38 genome to identify potential G4 regions based on the canonical G4 motif (G3+, N1−7, G3+, N1−7, G3+, N1−7, G3+). Bed files for individual chromosomes (1-22 & X) were created to have non-overlapping 100bp windows using bedtools and imported to R. Sequences were obtained for each bin using the Bioconductor package Biostrings (v2.62.2; BSgenome.Hsapiens.UCSC.hg38_1.4.4) and regions with missing base pairs (i.e. N) were removed. Bins were scored for G4 probability by DeepG4 and all bins with a probably score ≥ .99 were retained. Reads we counted over retained bins and compared across genotypes as described above in the regional analysis. Additionally, DeepG4 was used to scan enriched bins identified in the genome-wide relative enrichment analysis.

### Comparison of OG enrichment and RNA expression

To consider the impact of OG on RNA expression, data was compared between RNA-sequencing counts and OG-sequencing reads in promoter regions (defined at 1000bp upstream of the transcription start site (TSS)). OG-seq reads were counted over promoter regions and SES normalized input counts were subtracted from the OG signal. Counts were collapsed to meta-features by mean and normalized. RNA counts were mapped to OG reads by gene. Counts were averaged over replicates and genes were filtered if counts were not present in at least three of four genotypes per sequencing approach. Counts were z-scored by row within each sequencing approach and hierarchical clustering was performed using Euclidean distance to identify clusters that reliably showed a positive or negative correlation between OG and RNA across all genotypes (i.e. genes that show a consistent relationship of differing from the mean between oxidative damage and gene expression regardless of DNA glycosylase activity). To correlate changes in OG accumulation to changes in gene expression at specific genomic features, log2 fold changes for OG-DERs over the genomic feature and the corresponding log2 fold changes for the annotated genes from the RNA-seq analysis were extracted. Hierarchical clustering with a Pearson’s correlation coefficient and average linkage method was applied.

## RESULTS

### Loss of OGG1 and MUTYH impairs proliferation capacity and OG repair

HAP1 cells deficient for either MUTYH (*MUTYH*^-/^) or OGG1 (*OGG1*^-/^) or both DNA glycosylases (DKO) were generated by Horizon Discovery using CRISPR-Cas9 genome editing. The used guide RNA, targeted exon and insertion/deletion length is shown in Supplementary Table 1. Two independent clones of each knockout and wildtype (WT) cells were used for the experiments. To characterize the impact of DNA glycosylase deficiency, proliferation capacity of the cells and response to oxidative stress was monitored (Figure 1). Cell proliferation steadily increased over 72 h for all cell lines, but DKO showed a significantly reduced proliferation rate compared to WT from 48 h onwards (Figure 1A). Treatment with increasing concentration of hydrogen peroxide (H_2_O_2_) decreased cell survival accordingly but was not further affected in the knockout cells (Figure 1B). Additionally, we measured global OG levels by LCMS before and after treatment with 100 µM H_2_O_2_ (Figure 1C). No differences in non-treated conditions were found in single DNA glycosylase deficient cells compared to WT, but DKO showed significantly higher levels of global OG levels 24 h after treatment indicating impaired DNA repair.

### Fragmentation of DNA by sonication induces artificial OG

One crucial step during OG-seq is the initial DNA fragmentation, which determines the sequencing resolution. Fragments should be as small as possible to ensure a high resolution while still long enough to enable sequencing, ideally 150bp. Previously published OG-seq approaches employed sonication (18,20). However, sonication is a harsh treatment, with the potential of introducing artificial DNA oxidation. Here, we used a commercial nuclease mixture, Fragmentase®, to fragment genomic DNA and compared it to two sonication methods to investigate the effect on fragment shearing size and OG level increase. The Fragmentase® enzyme mixture digests DNA in a non-sequence specific manner and results in fragment sizes similar to sonication (Figure 2A lower panel). To identify the amount of artificial DNA oxidation introduced during sample preparation, we measured global level of OG by LCMS and found a wide variation among the three tested fragmentation methods (Figure 2A upper panel). Surprisingly, sonication increased global OG levels by 10-fold using the Covaris instrument and significantly by 39-fold for the Diagenode instruments compared to control. Enzymatically fragmented DNA did not significantly increase global OG levels (2.4-fold). Thus, DNA fragmentation using Fragmentase® minimizes artificial DNA oxidation providing a considerable improvement to the OG-seq pipeline.

**Figure 2:**
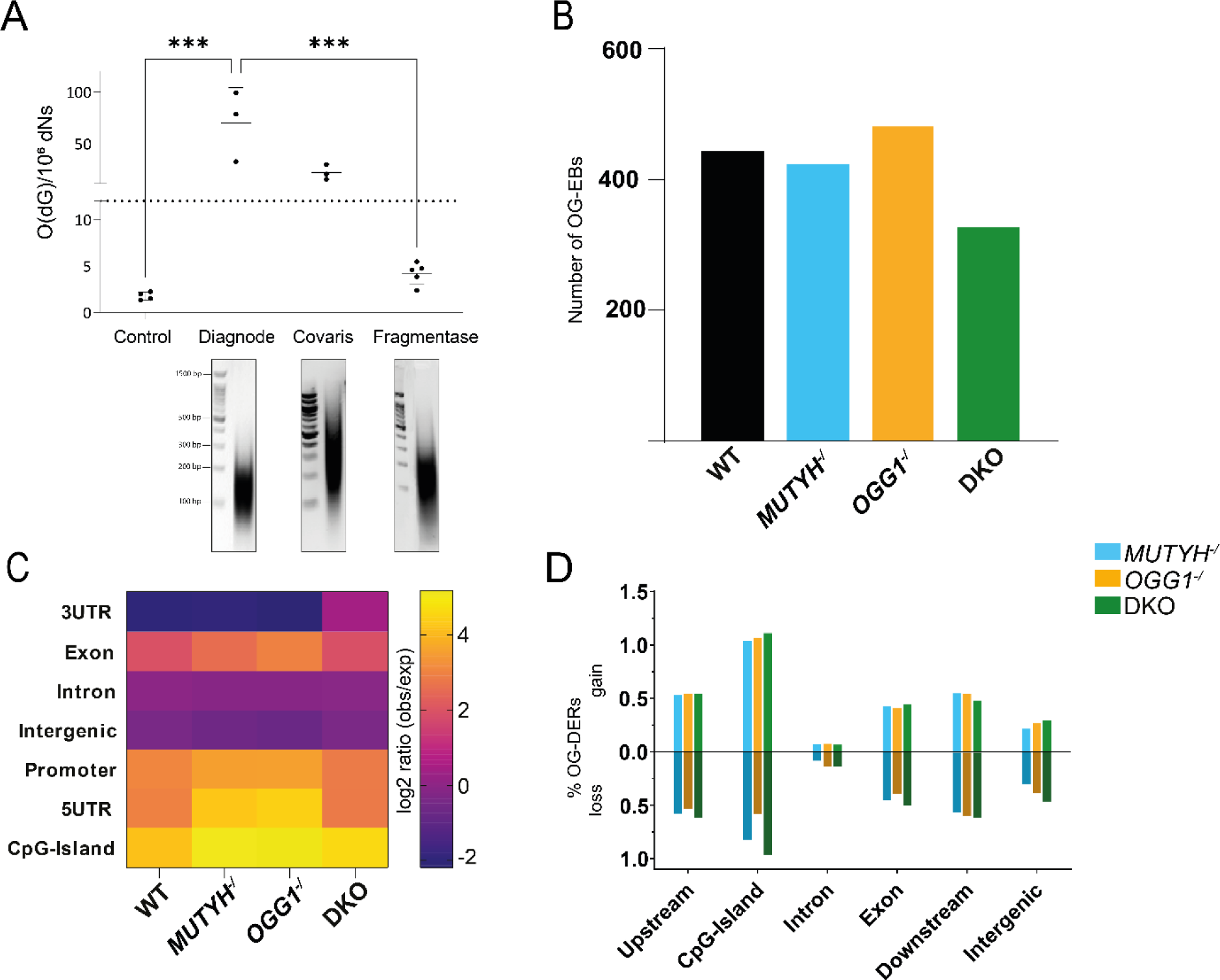
Optimization of fragmentation and OG-seq in WT and DNA glycosylase deficient HAP1 cells. **(A)** Comparison of different fragmentation methods. Top: LCMS was used to measure global O(dG) levels from Diagnode and Bioruptor sonicators as well as an enzymatic approach using Fragmentase. Error bars represent standard deviation from the mean, based on independent experiments (Control n=4, Diagnode n=3, Bioruptor n=3, Fragmentase n=5). Statistical analyses were done in GraphPad Prism software 10.1 using ordinary one-way ANOVA with Dunett’s multiple comparisons test. P-values < 0.05 were considered significant. Bottom: representative agarose gels from the respective fragmentation methods. **(B)** Number of OG enriched bins (OG-EBs) in WT, *MUTYH^-/^*, *OGG1^-/^*and DKO HAP1 cells. Log2 fold ratio of OG/input was calculated and enriched bins were defined as ≥ 95^th^ percentile within each genotype. **(C)** Heatmap of consensus peaks based on Homer annotation in studied genotypes, displayed as log2 ratio (observed/expected). **(D)** Percent of differentially OG-enriched regions (OG-DER) in different genomic features in *MUTYH^-/^*, *OGG1^-/^*and DKO compared to WT. Regions have been annotated to the nearest gene, among the different genomic features.

### OG is moderately enriched in the genome and predominantly located at CpG-rich regions

OG-seq, similar to immunoprecipitation-based sequencing methods (i.e. ChIP), involves identifying regions of read/signal enrichment in a targeted sample compared to a background sample. However, due to the ubiquitousness of OG across the genome, low frequency of occurrence and single-nucleotide nature of OG, classical peak-calling methods were unable to be applied to OG-seq data, a problem that has been reported previously (20). As such, a genome-wide relative enrichment analysis was used to identify OG-enriched bins (OG-EBs). OG-EBs had an up to 15% higher average GC content compared to the reported genomic GC content of 40.9% indicating specific OG pulldown (Supplementary Table 2). However, only a small number of OG-EBs in the genome were identified (average of 424 bins), with similar levels across the analyzed genotypes (Figure 2B). OG-EBs were annotated to their closest gene using Homer and found to be largely compromised of introns and intergenic regions (Supplementary Figure 1A). Using the ratio of observed to expected number of sequences, we found that our identified OG-EBs were overrepresented for regulatory genomic features including promoter regions, 5′ untranslated regions (UTR) and CpG-islands (Figure 2C). On the other hand, OG-EBs were underrepresented in 3′ UTR and intergenic regions. This correlated well with the CpG content of the genomic features, with an increased enrichment of OG-EBs in CpG-richer regions (Supplementary Table 3). OG-EBs were similarly distributed across genomic features between genotypes (Figure 2C), however, analysis of the annotated genes showed almost no overlap (Supplementary Figure 1B). This indicates that though OG frequency is related to the genomic landscape, it appears to be stochastic in relation to specific genes. Despite, GO pathway analysis of OG-EBs revealed similar pathways across genotypes which were predominately related to system and cellular development as well as cellular growth and differentiation (Supplementary excel file 1).

To identify DNA glycosylase specific differences, we compared OG accumulation between WT and knock out cells across specific genomic features. We found similar levels of gain or loss of OG within each genomic feature between all genotypes (Figure 2D). CpG-Islands showed the highest percentage of OG-differentially enriched regions (OG-DERs) affecting on average 1.86% of all CpG-islands analyzed. Introns were the least affected by loss of DNA glycosylases, with on average only 0.19% of these regions being identified as OG-DERs. Pathway analysis of OG-DERs across all the analyzed genomic locations and genotypes revealed a few significant hits which mainly related to processes with a broader biological role in development (Supplementary excel file 2). We further focused on OG-DERs found within genomic features that demonstrated most differences in OG enrichment including CpG-islands, upstream regions and exons. GO analysis of either gain or loss of OG in the KOs across all three regions identified no significant (FDR≤.05) pathways in *MUTYH^-/^*, but pathways for *OGG1^-/^* and DKO associated with chemical stimuli, biogenesis, and development (Supplementary excel file 2). Despite similar levels of loss and gain across our KOs, the overlap of OG-DERs annotated genes between the genotypes were substantially higher in genes with OG gain (69-73%) than OG loss (30-40%) across all three regions.

### OG accumulates in the context of G4-structures

Previous studies reported differences in the occurrence of OG within G4-structures (18,20,22), thus we investigated the distribution of predicted G4-forming sequences over OG-EBs in our cell lines. We found 0.05% of all analyzed bins across the genome contained high probability G4-forming motifs (≥.99). In contrast, we observed around 18% of the OG-EBs in our genotypes contained G4-forming motifs with the highest level seen in *OGG1*^-/^ (Figure 3A). To find OG-DERs within G4’s (G4-DERs), we first identified high probability G4-motifs across the human reference genome (hg38). By using 100bp bins and a probability score ≥.99, we identified 13 256 bins containing potential G4-forming sequences. These G4 regions demonstrated a high CpG content comparable to CpG islands. (Supplementary Table 3). We found that up to 2% of the G4-regions had different OG accumulation in DNA glycosylase deficient cells and most of these showed gain of OG (Figure 3B). However, G4-DERs showed similar levels of gain and loss across all genotypes when compared to WT. GO analysis of G4-DERs revealed pathways related to cellular differentiation, proliferation, homeostasis, and response to stimuli, however these did not maintain significance after FDR correction (Supplementary excel file 3).

**Figure 3:**
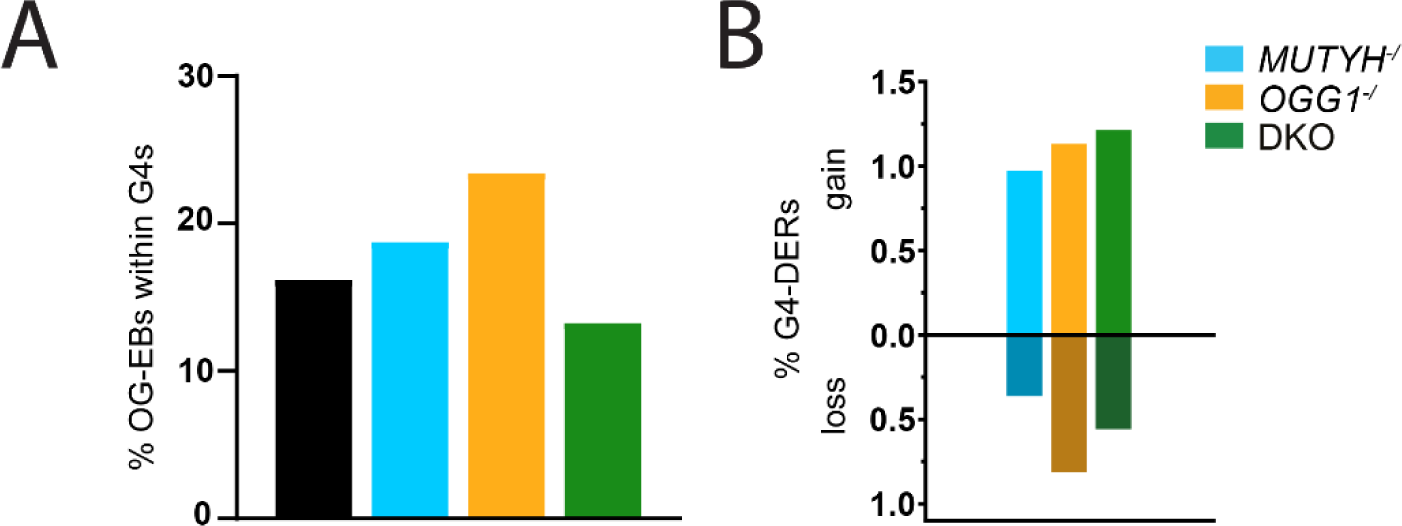
Enrichment of OG at G4-forming sequences in WT and DNA glycosylase deficient HAP1 cells. **(A)** Percent of OG enriched bins (OG-EBs) containing G4-forming sequences of WT, *MUTYH^-/^*, *OGG1^-/^* and DKO HAP1 cells. G4-forming sequences have been identified using DeepG4 with a probably score of ≥ .99. **(B)** Percent of differentially OG-enriched regions within G4 motif regions (G4-DER) in *MUTYH^-/^*, *OGG1^-/^* and DKO compared to WT.

### DNA Glycosylase-dependent changes in gene expression

Next, we performed RNA-seq to investigate the impact of DNA glycosylase-deficiency on gene expression. The lack of MUTYH caused 212 differentially expressed genes (DEGs), compared to WT, while the lack of OGG1 caused 299 DEGs. The absence of both DNA glycosylases resulted in 317 DEGs (Figure 4A). Overall, only 28 genes were found to be differentially expressed in all three genotypes, but between 40-46% of DEGs of each individual genotype were found in at least one other genotype (Supplementary Figure 2). When looking across all DEGs independent of genotype using GO, we found a broad spectrum of metabolic and developmental processes along with cell differentiation, gene expression and RNA processing (Figure 4B). When analyzing the DEGs found in at least two genotypes, specific developmental pathways (i.e. muscle, organ, circulatory development) as well as response to oxygen-containing compound were overrepresented (Supplementary excel file 4).

**Figure 4:**
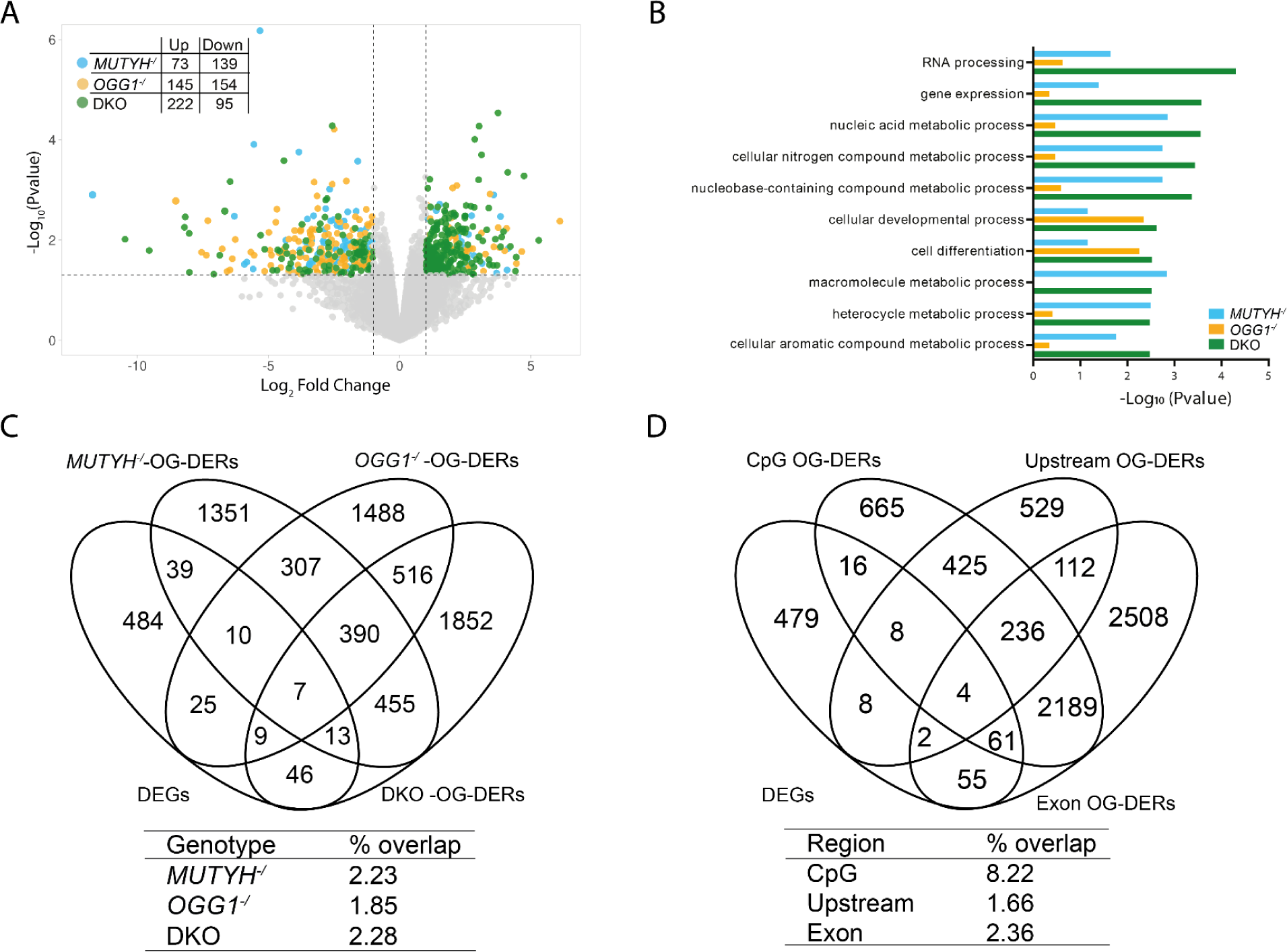
Correlation of RNA-seq with OG-seq in WT and DNA glycosylase deficient HAP1 cells. **(A)** Volcano plot of up and down regulated genes identified by RNA-sequencing from *MUTYH^-/^*, *OGG1^-/^*and DKO HAP1 cells compared to WT. Dotted lines represents FC >±1 (log2) and p-value <0.5 (-log10) **(B)** Top ten enriched Gene Ontology spathways ranked by p-value and identified by protein analysis through evolutionary relationships (PANTHER). **(C)** Overlap of all differentially expressed genes (DEGs) from RNA-seq with OG differentially enriched regions (OG-DERs) from *MUTYH^-/^*, *OGG1*^-/^ and DKO across exons, CpG islands and upstream region. (D) Overlap of all DEGs from RNA-seq and OG-DERs between exons, CpG islands and upstream regions across all genotypes.

### Stress response genes regulate their transcription dependent on OG

To address the epigenetic-like function of OG, genome-wide OG distribution was correlated with the gene expression profile of all genotypes. First, we compared the DEGs across all genotypes (n=633) to OG-EBs that were found in the KOs but not in the WT and found little overlap (n=38). Similarly, the overlap of G4-DERs (n=471) and DEGs resulted in little overlap (n=8). Next, we focused our analysis on the overlap of OG-DERs from CpG-islands, upstream regions, and exons with DEGs. Venn analysis by genotype showed that about 2% of the OG-DERs from any one genotype could be found amongst DEGs, but there was no difference in overlap rates between genotypes (Figure 4C). Venn analysis by genomic feature across genotypes showed that OG-DERs in exons and upstream regions had a small overlap with DEGs of 2.4% and 1.7% respectively (Figure 4D). In contrast, OG-DERs from CpG islands showed an 8.2% overlap with DEGs. Pathway analysis revealed that overlapping genes were related to developmental pathways (Supplementary excel file 5). The comparison of RNA-seq reads and OG-seq reads from overlapping genes in the analyzed regions revealed both a negative and positive effect of OG on gene expression (Supplementary Fig 3).

To further investigate the correlation between OG accumulation and transcription, we compared all RNA-seq reads and OG-seq reads in gene promoters (defined as 1000bp upstream of the TSS). Using hierarchical clustering of scaled read counts we focused on genes that showed consistent positive or negative correlation between OG levels and RNA expression regardless of genotype. Using GO pathway analysis, we found a significant overrepresentation of genes related to metabolic processes amongst positively correlated genes, while genes that consistently showed a negative relationship between OG- and RNA-seq had an overrepresentation of genes related to cellular (re)organization, DNA damage response and cell cycle (Table 1). These results indicate a DNA glycosylase-independent effect on transcription by OG at promoters specifically of genes related to the stress-response systems.

**Table 1:** Top 5 highest ranked GO pathways from either positively or negatively correlated OG-seq and RNA-seq read counts at promoter regions (defined as 1000bp upstream of the transcription start site).

| **GO biological process** | **Fold enrichment** | **FDR** |
| --- | --- | --- |
| **positive correlation** |  |  |
| cellular metabolic process | 1.16 | 0.00669 |
| primary metabolic process | 1.14 | 0.00302 |
| nitrogen compound metabolic process | 1.14 | 0.00850 |
| metabolic process | 1.13 | 0.00311 |
| organic substance metabolic process | 1.13 | 0.00423 |
| **negative correlation** |  |  |
| vacuole organization | 2.14 | 0.03530 |
| DNA damage response | 1.63 | 0.00188 |
| cell cycle | 1.44 | 0.00695 |
| organelle organization | 1.30 | 0.00060 |
| cellular localization | 1.25 | 0.00416 |

## DISCUSSION

Oxidative DNA damage and its repair has been shown to be involved in transcriptional regulation but whether these processes are directly linked remained largely understudied. Correlating genome-wide OG distribution and the gene expression profile of OGG1 and/or MUTYH-deficient cells, we present evidence that both OG and DNA glycosylases modulate gene transcription independently.

OGG1 and MUTYH prevent the mutagenic effect of OG and synergistically promote survival in mice (29). Using HAP1 cells deficient for both DNA glycosylases, we observed decreased cell proliferation after longer culturing periods but no additional effect on cell survival after oxidant treatment. This is partially in contrast with a previous study, using primary fibroblast from DKO mice that also found slower proliferation but increased sensitivity to H_2_O_2_ treatment (30). However, OGG1 and/or MUTYH-deficient HAP1 cancer cells may have developed strategies to adjust to increased oxidative stress and maintain resistance to apoptosis which has recently been demonstrated for NEIL DNA glycosylases (31).

Despite lacking DNA glycosylases, we found no significant global accumulation of OG in untreated cells compared to WT. Previous studies of OGG1 and/or MUTYH deficient models reported, dependent on tissue, age, and oxidative stress exposure, both no change in OG accumulation as well as significantly increased OG. In the liver, *Ogg1^-/-^* and DKO showed higher OG accumulation that continued to increase with age, while no differences were found in kidney or spleen (32–34). In the brain, no significant differences in OG accumulation have been found in Ogg1, Mutyh and DKO mice (3,34). To our knowledge, no other study has quantified OG in DNA glycosylase deficient HAP1 cells, however, one study analyzed repair capacity and found that OGG1 deficient HAP1 cells showed a higher rate of BER than WT cells at baseline (35) suggesting efficient compensation of other available DNA glycosylases in the absence of OGG1. A potential candidate is Nth-like DNA glycosylase 1 (NTHL1) that was shown to repair OG (36) and is also expressed at high levels in HAP1 cells (31). Other studies demonstrated that NEIL1 can act on OG and may also serve as a potential backup to OGG1 (37,38). We speculate that each DNA glycosylase shows a high affinity for a specific type of DNA lesion and lower affinity towards other lesions, thus when higher affinity proteins (i.e. OGG1 to OG) are available they will preferentially bind and repair. In the absence of high affinity competition, proteins with lower affinity (i.e. NTHL1 and NEIL1) can bind to compensate for the loss. Additionally, Jun et al. (35) found that when treated with an oxidative agent compensatory repair of OG was sufficient during acute exposure (2 h) but was unable to cope efficiently during prolonged exposure (24 h). In line with this, we found a significant increase of OG in our DKO cells 24 h after acute oxidative exposure. This compounded effect from the loss of MUTYH in addition to OGG1 suggests that, like OGG1, compensatory glycosylases are unable to excise OG when it has already mispaired (39).

Given the proposed epigenetic role of OG, sequencing its genomic distribution is of high interest. However, background oxidation of DNA during sample preparation can artificially increase OG and compromise data interpretations. Here, we identified sonication as a major source of artificial OG introduction which may explain previous results that reported several thousands of OG enriched regions as compared to our 448 OG-EBs in WT HAP1 cells using our improved approach (17,18). To our knowledge, only one study analyzed genome-wide OG distribution in OGG1-deficient cells and found a 2-fold increase in peaks of OG enrichment (18). However, these results were based on one sequencing sample prepared using sonication and should be taken with caution. We found no major difference in OG enrichment after loss of OGG1 and/or MUTYH which is in line with the similar global levels of OG and may be explained by compensatory repair mechanisms as discussed before. In summary, by minimizing the presence of artificial OG using enzymatic digestion, one can ensure that the level of OG throughout the genome is better reflecting the *in vivo* situation and conclusions drawn are biologically relevant.

Most of our OG-EBs were found in introns and intergenic regions as described previously (20). However, after accounting for the relative size of genomic features across the genome, we found an overrepresentation of enriched bins in gene regulatory regions like promoters, 5′ UTRs and CpG islands, in line with what has been reported by others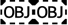. Given the higher G content of CpG-islands the likelihood of OG accumulation is consequently higher. A similar bias has been described before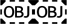. In contrast, sequencing of OG in single base resolution suggested a bimodal model for OG accumulation at high CpG promoters with a 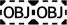 which will be missed with our OG-seq approach because of the lower resolution.

The overall level of OG accumulation in specific genomic features did not differ between KOs and WT, only the precise location of OG. Moreover, the number of OG-DERs were generally low suggesting that OG distribution is not notably impacted by its repair. However, some genomic features showed a greater difference in OG accumulation in KOs, and this correlated directly with the GC% in those regions, with CpG-islands showing the highest percentage of OG-DERs and highest GC content. Together, this indicates that OG accumulates randomly within a genomic feature and differences are mainly driven by region rather than genotype. This is further underlined by pathway analysis which revealed no significant overrepresentation of pathways in all regions and all genotypes with few exceptions. However, in a previous study (19), a catalytic inactive OGG1 was found bound to OG mainly at the same regions that we identified as differentially enriched in the KOs (e.g. upstream and CpG islands) suggesting that at least partially OG might be preferentially repaired at gene regulatory regions by OGG1.

Another genomic feature with higher G:C content is G4 structures, which are found at promoter regions (40). In line with previous studies, we identified an enrichment of OG at G4 sites (19,20) which further increased upon loss of OGG1 (18). More recently, however, it was shown that OG is depleted at G4 structures that have been detected in living cells (22). Since G4 distribution will depend on cell type and to our knowledge no G4 profiling in HAP1 cells has been performed, we cannot exclude that OG-EBs within G4 sequences are only G4s in their unfolded, duplex DNA state. The different knockout models showed minor differences in relative G4 enrichment suggesting that OG distribution at G4 sites is mainly independent of its repair. Pathway analysis of G4-DERs compared to all G4 regions did not find any significant overrepresentation of pathways neither within nor across genotypes, further highlighting that OG accumulation within G4 regions seems to occur randomly. One study mapped the binding of acetylated OGG1 in human cells and found a positive correlation with genome-wide occurrence of G4s and reduced G4 foci in OGG1-deficient MEF cells suggesting that OGG1-initiated repair is important for G4 formation (41). However, direct evidence is missing, and more studies are needed to fully understand the link between OG and DNA glycosylases at G4 sites.

We have previously shown that loss of OGG1 and/or MUTYH in mice impact gene transcription independent of global accumulation of OG (3). However, a contribution of gene region-specific OG accumulation driving gene expression changes could not be excluded. Here, we found no correlation between OG-DERs and gene expression changes, and a minor overlap of OG-DERs with DEGs in the knockout cells. This demonstrates that OGG1 and MUTYH can modulate gene transcription independent of OG repair. In support of our finding, a previous study demonstrated that catalytic inactive OGG1 could facilitate pro-inflammatory gene expression (42). However, we cannot exclude that under oxidative stress conditions DNA repair-dependent mechanisms occur as proposed by others (43–45). The underlying mechanism of DNA glycosylase repair activity-independent gene regulation is still unknown. Several studies suggest them as modifiers of the epigenetic landscape (46–49) and demonstrate their interaction with epigenetic modifiers such as DNA methyltransferases (50), histone demethylases (51) and members of the polycomb repressor complex (PRC) (52). Interestingly, DEGs of our knockout cells showed highest enrichment in target genes of PRC2 including SUZ12 (adjusted p-value: 4.690e-18), JARID2 (adjusted p-value: 8.007e-15) and EZH2 (adjusted p-value: 4.194e-11) as identified by Enrichr (53). Our group previously found that DEGs from the hippocampus of Ogg1- and/or Mutyh-deficient mice significantly overlapped with genes bearing histone trimethylation at K27 (3), an epigenetic mark which is placed by PRC2 (54). Together, this suggests that OGG1 and MUTYH may cooperate with PRC2 to modulate the epigenome and hence gene transcription.

Nevertheless, several studies suggest that OG itself can regulate gene expression particularly when located in promoter regions (43,55). Thus, we filtered our RNA-seq dataset on genes that showed a consistent relationship between OG accumulation in promoters and gene expression regardless of DNA glycosylase activity and show that OG can drive both transcriptional repression and activation, particularly of genes involved in general stress response pathways. This suggests that the accumulation of OG may serve as an oxidative stress sensor for those genes triggering cellular responses, whereby the cell works to meet the energy and biosynthetic demands caused by the stress condition (56). This impact of OG on transcription, however, is independent of its repair and might instead be facilitated by physically blocking the binding of transcription factors (57,58) or altering the formation of secondary DNA structures (59,60).

In conclusion, our findings reveal distinct pathways of transcriptional regulation by OG and its repair proteins OGG1 and MUTYH. Whereas OG accumulation is driven by the sequence context to signal cellular stress, OGG1 and MUTYH may have more specialized functions possibly as epigenetic modifiers.

## DATA AVAILABILITY

All genomic data produced in the present project (OG-seq and RNA-seq) have been deposited in the NCBI GEO database under accession numbers GSE256080 and GSE256081, respectively.

## SUPPLEMENTARY DATA

Supplementary data are available Online.

## ACKNOWLEDGEMENTS

We thank Christine Gran Neurauter for verifying the genotype of the DNA glycosylase-deficient HAP1 clones and the Proteomics and Modomics Experimental Core Facility (PROMEC) at NTNU for the help with mass spectrometry analysis.

