## Supplementary Data for "Genomic 8-oxoguanine modulates gene transcription independent of its repair by DNA glycosylases OGG1 and MUTYH"

Supplementary Table 1 - 3

Supplementary Figure 1 -3

**Supplementary Table 1: CRISPR/Cas9-induced InDels of *OGG1* and *MUTYH*.**

The length of the insertion (In) or deletion (Del) is indicated.

| Target gene | Clone | Parental clone | Target exon | Guide RNA | InDel |
|---|---|---|---|---|---|
| *OGG1* | 1 | WT HAP1 | 1 | GTACGATGCCCCATGCGCCT | 5 bp Del |
| | 2 | WT HAP1 | 1 | GTACGATGCCCCATGCGCCT | 8 bp Del |
| *MUTYH* | 1 | WT HAP1 | 3 | CTTGGTCGTACCAGCTTAGC | 13 bp Del |
| | 2 | WT HAP1 | 3 | CTTGGTCGTACCAGCTTAGC | 5 bp Del |
| *OGG1* and *MUTYH* (DKO) | 1 | OGG1 clone 2 | 3 | CTTGGTCGTACCAGCTTAGC | 1 bp Ins |
| | 2 | OGG1 clone 2 | 3 | CTTGGTCGTACCAGCTTAGC | 5 bp Del |

**Supplementary Table 2: Reported and observed global GC content of OG enriched regions from OG-seq**

| | **Average GC-content** |
|---|---|
| Reported global | 40.9 % |
| WT | 49.9 % |
| *MUTYH*$^{-/-}$ | 53.5 % |
| *OGG1*$^{-/-}$ | 56.3 % |
| DKO | 50.1 % |

**Supplementary Table 3: GC content of different genomic features used in regional analysis**

| Region | Average GC-content |
|---|---|
| Intergenic | 46.8% |
| Upstream | 48.3% |
| Introns | 45.9% |
| Downstream | 43.8% |
| CpG island | 68.7% |
| Exons | 50.8% |
| G4 | 63.0% |

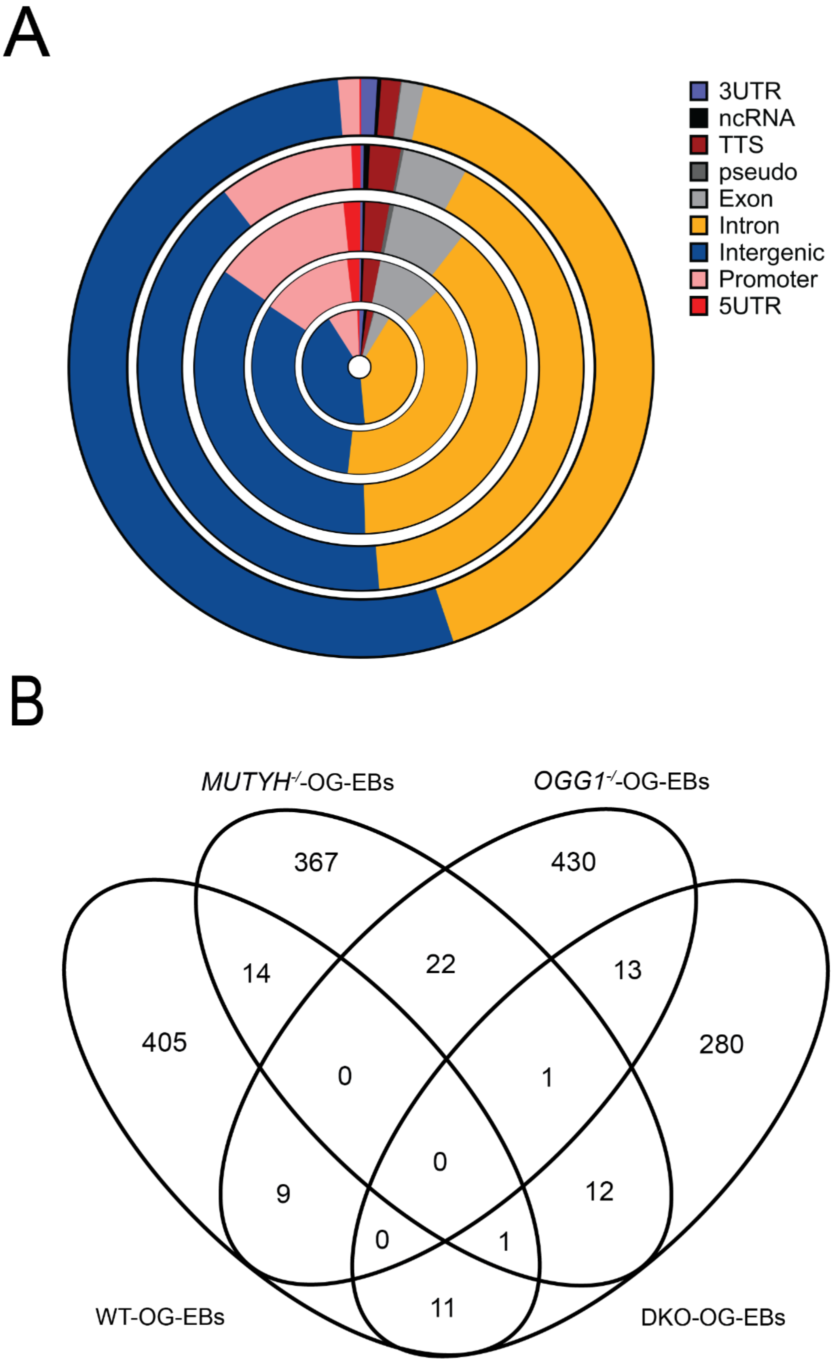


Supplementary Figure 1: Genomic distribution of OG enriched bins and overlap between genotypes. (A) Distribution of OG-enriched bins (OG-EBs) in different genomic features. Outer Ring towards inner ring: expected over the whole genome, WT, *MUTYH*$^{-/}$, *OGG1*$^{-/}$ and DKO. (B) Overlap of annotated genes in OG-EBs across the genotypes.

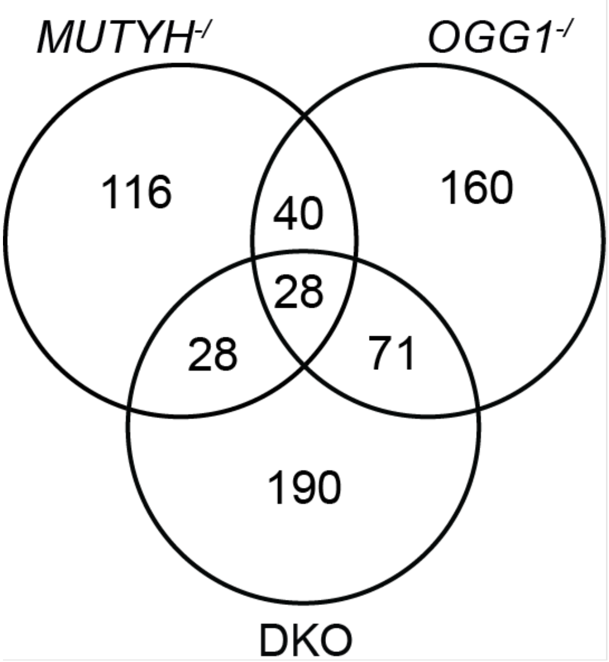


Supplementary Figure 2: Overlap of differentially expressed genes in *MUTYH*$^{-/}$, *OGG1*$^{-/}$ and DKO as identified by RNA-seq.

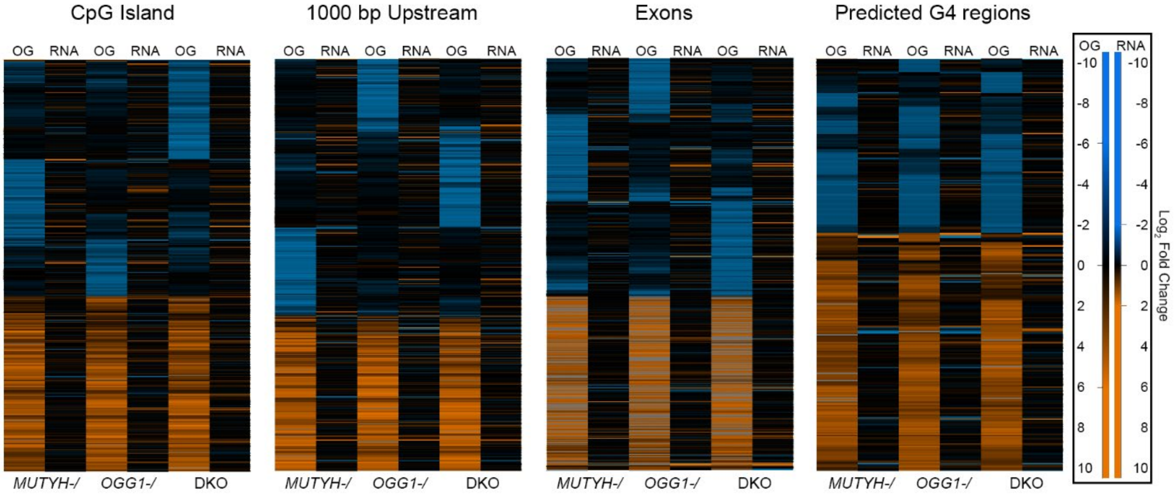


Supplementary Figure 3: Comparison of RNA-seq and OG-seq in different genomic features across DNA glycosylase deficient cells. Clustered heatmaps showing $\log_2$ fold change of differentially enriched regions (OG-DERs) in OG sequencing and corresponding $\log_2$ fold change of RNA sequencing across *MUTYH-/*, *OGG1-/* and DKO HAP1 cells. Blue indicates lower OG accumulation and lower gene expression compared to the WT control, while orange indicates increased OG accumulation or gene expression.
